# VRK1 kinase maintains an undifferentiated proliferative state in neuroblastoma tumor cells

**DOI:** 10.64898/2026.08.05.742965

**Authors:** Mónica Ojeda-Puertas, María A. Gómez-Muñoz, Ana Colmenero-Repiso, Aida Amador-Álvarez, Ismael Rodríguez-Prieto, Ricardo Pardal, Francisco M. Vega

**Affiliations:** Instituto de Biomedicina de Sevilla (IBiS) (Hospital Universitario Virgen del Rocío/CSIC/Universidad de Sevilla); Departamento de Biología celular, Facultad de Biología, Universidad de Sevilla, 41012 Seville, Spain; Departamento de Fisiología Médica y Biofísica, Universidad de Sevilla, 41013 Seville, Spain; Department of Pathology, Smilow Research Center, NYU Grossman School of Medicine, New York 10016, USA

**Keywords:** neuroblastoma, high-risk, VRK1, SOX2, differentiation, stemness, cancer stem cells, neural crest

## Abstract

Neuroblastoma is a neural crest-derived pediatric malignancy characterized by marked cellular heterogeneity and variable differentiation status. Undifferentiated tumors are associated with aggressive clinical behavior, treatment resistance and poor outcome, highlighting the need to identify molecular mechanisms that sustain tumor cell plasticity and prevent differentiation. Vaccinia-related kinase 1 (VRK1) is a serine/threonine kinase involved in cell-cycle progression, DNA-damage responses and transcriptional regulation, and has previously been associated with neuroblastoma progression. However, its role in the control of neuroblastoma differentiation remains unclear.

Here, we investigated the relationship between VRK1 expression, tumor differentiation and stem-like properties in human neuroblastoma. Analysis of patient tumor datasets and tissue microarrays showed that VRK1 expression is enriched in undifferentiated neuroblastoma and stage 4 tumors, and inversely correlates with established differentiation markers, including DDC, NCAM1 and S100B. This association was maintained in MYCN-non-amplified tumors, indicating that the relationship between VRK1 and differentiation is not dependent on MYCN status. Single-cell transcriptomic analyses further demonstrated elevated VRK1 expression in developmentally immature neural crest progenitor and Schwann cell precursor-like populations.

Induction of neuronal or mesenchymal differentiation consistently reduced VRK1 expression in neuroblastoma cell lines and patient-derived cells. Conversely, VRK1 silencing promoted differentiation-marker expression, reduced nestin and Ki67 expression, and produced sustained differentiation-associated changes in xenograft tumors. VRK1 was also enriched in tumorsphere cultures that select for undifferentiated stem-like neuroblastoma cells. VRK1 depletion impaired tumorsphere growth, reduced intratumoral proliferation and altered the balance between undifferentiated cells and differentiated progeny, supporting a role for VRK1 in self-renewal and maintenance of progenitor-like tumor cells. Mechanistically, VRK1 expression positively correlated with the core stemness transcription factor SOX2 in neuroblastoma tumor cells. VRK1 knockdown reduced nuclear SOX2 abundance, whereas VRK1 overexpression increased SOX2 protein levels. In addition, analysis of the VRK1 locus identified an active chromatin configuration and potential SOX2-binding sites, consistent with a regulatory relationship between these factors. Together, these findings identify VRK1 as a regulator of the undifferentiated, proliferative and stem-like state in neuroblastoma. The VRK1-SOX2 axis may contribute to stabilizing tumor-cell immaturity and represents a potential target for differentiation based therapeutic strategies in high-risk neuroblastoma.

## Background

Neuroblastoma (NB) is the most common cancer diagnosed during the first year of life and the most frequent extracranial pediatric solid tumor (1). It arises during embryonic development from sympathoadrenal precursors of the neural crest (2). Neuroblastoma is characterized by a remarkable clinical and biological heterogeneity, with patients with aggressive forms of the disease frequently presenting extensive metastasis and relapse, and exhibiting event free survival rates below 50% (3). A key determinant of neuroblastoma behavior is the degree of tumor differentiation, with undifferentiated tumors associating with advanced stage, poor prognosis, and resistance to conventional therapies (4). The study of NB cellular heterogeneity and plasticity by single cell transcriptomics and other technologies has demonstrated the presence of undifferentiated, persistent, progenitor-like cell populations associated with relapse and aggressiveness in high-risk NB (5–7). Understanding the molecular mechanisms that control neuroblastoma differentiation is crucial for developing therapeutic strategies that can redirect tumor cells toward less aggressive, differentiated phenotypes.

VRK1 (Vaccinia-related kinase 1) is a serine/threonine kinase that plays critical roles in cell cycle progression, DNA repair, and transcriptional regulation (8–10). VRK1 is highly expressed in proliferating tissues and has been implicated in various cancers, where it typically promotes cell proliferation and survival (10–17). In neuroblastoma, we previously described that VRK1 is highly expressed in proliferating neuroblasts, being a marker for tumor progression and malignancy, independently of MYCN amplification (18).

Studies have suggested a role for VRK1 in stem cell biology and during development, with evidence indicating that VRK1 may help maintain pluripotency and prevent differentiation in embryonic stem cells (19). VRK1 expression peaks at the transit amplifying undifferentiated compartment in various regenerative tissues (19), is a main player during embryonic development of hematopoiesis (20) and is essential for spermatogonia cell maintenance (21,22).

We previously found that a VRK1-associated expression signature is significantly associated with signaling pathways regulating proliferation and differentiation in neuroblastoma tumors (18). Given VRK1’s established functions in cell cycle control and its emerging roles in stemness maintenance, we hypothesized that VRK1 might contribute to maintaining neuroblastoma cells in an undifferentiated, proliferative state and could represent a novel target for differentiation therapy. In this study, we investigated VRK1 expression patterns in human neuroblastoma samples and examined its functional role in controlling differentiation using several in vitro models. Our findings reveal that VRK1 is preferentially expressed in undifferentiated neuroblastoma and that functions as a key regulator preventing tumor cell differentiation through modulation of the core stemness transcription factor SOX2.

## Methods

### Transcriptomic data analysis in neuroblastoma patient tumors

Gene expression data from neuroblastoma patient cohorts were obtained and analyzed from public databases included in R2 Genomics Analysis and Visualization Platform (http://r2.amc.nl). Databases used were GSE45547, GSE49710 and GSE3446. The data were managed and processed with Excel and Prism and displayed via box plots, violin plots or heatmaps. Kaplan‒Meier survival analyses were performed via the R2 platform by Kaplan-scan method with Bonferroni-corrected p-values. VRK1 expression levels were correlated with differentiation markers (DDC, NCAM1, S100B) and tumor stage using Pearson correlation analysis.

For functional enrichment analysis, tumors (GSE49710) with a high expression of VRK1 were stratified according to histology (favorable versus unfavorable), and the genes correlating with a high expression of VRK1 (r>0.5; p-value<0.05) in each group were identified. The resulting 3 list of genes (high in favorable only (374 genes), high in unfavorable only (727 genes) and high in both (85 genes)) were analyzed in Enrich (https://maayanlab.cloud/Enrichr) against the following databases: MSigDB Hallmarks v2023.2, KEGG 2021 Human, Reactome 2022, GO Biological Process 2023, MSigDB Oncogenic Signatures, CellMarker 2024, JASPAR 2022 and MSigDB Computational. For each database, the adjusted p-value, odds ratio and combined score were obtained. Normalized enrichment scores across relevant functional axes were represented. Scores were calculated by normalizing the −log_10_(adjusted p-value) of the most significant term per category against the global maximum observed (E2F Targets, Common group = 109.2).

Single cell transcriptomic visualizations were obtained by using the DotPlot and VlnPlot functions of the Seurat R package, utilizing publicly available scRNA-seq data.

For VRK1 transcription regulation analysis, the VRK1 gene structure (chr14:96,797,382–96,881,609; GRCh38/hg38, NCBI Gene ID 7443) was retrieved from NCBI (https://www.ncbi.nlm.nih.gov). Published SOX2 ChIP-seq peaks were obtained from the non-redundant ReMap2022 atlas (remap2022_SOX2_nr_macs2_hg38_v1_0) and filtered for chr14 peaks. Peaks were annotated by contributing cell-type biotype. Gene SOX2 binding motifs were predicted using the JASPAR 2024 CORE vertebrate matrix MA0143.4 (relative score threshold ≥0.80) across six genomic windows: the proximal promoter (chr14:96,792,382–96,798,382) and five additional intragenic/downstream regions previously identified from ReMap peak coordinates (chr14:96,820,000–96,824,000; 96,862,000–96,866,000; 96,877,000–96,881,000; 96,880,000– 96,884,000; 96,887,000–96,891,000). For each region, motif genomic coordinates were reconstructed by adding the FASTA sequence offset to the local motif start/end positions reported by JASPAR. Motifs with relative score ≥0.90 were classified as high confidence. Motif-peak overlap was determined by direct genomic coordinate intersection between JASPAR motif boundaries and ReMap peak boundaries. NarrowPeak ChIP-seq peak calls for H3K4me3 and H3K27ac were obtained from the ENCODE portal (https://www.encodeproject.org/) for SK-N-SH neuroblastoma cells, human neural crest cells, and human neural stem cells. All genomic track visualizations were generated in Python using Plotly.

### Tissue analysis by immunohistochemistry

Human neuroblastoma tissue samples (n=31) were obtained from the institutional biobank, with appropriate ethical approval, and placed on a tissue microarray (TMA) (18). Tumors on the TMA were clinically classified as differentiated or undifferentiated based on histological examination. Immunohistochemistry for VRK1 was performed using standard protocols. Briefly, paraffin-embedded tissue sections were hydrated, subjected to antigen retrieval with citrate buffer (pH 6), and incubated with the following primary antibodies: rabbit anti enolase (Sigma, San Luis, MO, USA; 1:1000), DDC rabbit (1:1000, Millipore), Ki67 rabbit (1:200, Thermo Scientific), and rabbit anti VRK1 (Sigma, San Luis, MO, USA; 1:500). Secondary antibodies used were biotinylated-conjugated anti-mouse and biotinylated-conjugated anti-rabbit (Vector Laboratories Burlingame, CA, USA; 1:1000). The Vectastain ABC kit and DAB peroxidase substrate (Vector Laboratories) were used for staining. Images were analyzed with QuPath software (23). Pathologists scored the samples semi-quantitatively using a scale that combines the percentage of positive cells and the intensity of the reaction product (24).

### Cell culture and transfection

The human NB cell lines SK-N-SH and IMR-32 were obtained from the American Type Culture Collection (ATCC) and cultured in DMEM + GlutaMAX (GIBCO BRL) supplemented with 10% fetal bovine serum (FBS) (GIBCO BRL), 100 µg/ml streptomycin, and 100 U/ml penicillin. The NB48T PDX primary cell line was derived from freshly obtained tumors and have been described previously (25). All cells were maintained at 37°C in a 5% CO2 humid atmosphere.

For transfection, cells at 80-90% confluence were transfected with 2.5-5 µg of plasmid DNA (pCEFL-KZ-HA-VRK1) or 50-100 nM siRNA against VRK1 (siVRK1-02: 5’-GGUGGAACUUCGAGAAGAA-3’ and siVRK1-03: 5’-GGACAAGGUCUUCGAGAUU-3’, Horizon) or non-targeting siRNA control (siControl, Horizon) using Lipofectamine 2000 (Life Technologies) according to manufacturer’s instructions. Knockdown efficiency was confirmed 48-72 hours post-transfection.

For differentiation assays, NB cells were seeded in complete medium and treated 24h after with standard culture media or the same media containing 10 µM all-trans retinoic acid (ATRA) or 5 ng/ml TGF β1 (R&D) for 7-10 days, as described in (25).

### Western Blot Analysis

Protein lysates were prepared using lysis buffer (50 mM Tris-HCl (pH 8), 0.5 mM EDTA, 150 mM NaCl, 1% Triton X-100, protease, and phosphatase inhibitor cocktails). Equal amounts of protein (30-50 μg) were separated by 10% SDS-PAGE and transferred to PVDF membranes (Millipore). Primary antibodies against VRK1 (1:1000, Sigma-Aldrich), Nestin (1:500, Millipore), NCAM1 (1:1000, Millipore), SOX2 (1:1000, Santa Cruz Biotech.), and GAPDH (1:2000, Trevigen) were used. Protein bands were visualized using HRP secondary antibodies (1:1000, Jackson ImmunoResearch) and ECL detection reagent (GE Healthcare) in a ChemiDoc™ Touch Imaging System (BioRad) and quantified using FIJI software (NIH).

### Tumorsphere formation assay

For undifferentiated tumorsphere formation, SK-N-SH cells were serially cultured in neural crest media (DMEMF-12 medium supplemented with 1% N2, 0.5% B27, 15% FBS, 100 µg/ml streptomycin, 100 U/ml penicillin, 10 μg/ml FGF, 20 μg/ml IGF-1, and 20 μg/ml human EGF) at low density in low-adherence plates as previously described (26). Briefly, spheres were allowed to form for 7-10 days at 37°C, then counted, cryopreserved and processed for immunofluorescence. For gene expression analysis, spheres were collected and processed for RNA extraction. When indicated, cells were transfected with the corresponding siRNA using reverse transfection in suspension cell cultures before tumorsphere formation assay.

### Immunofluorescence

For immunofluorescence, cells were fixed with 4% PFA in PBS for 15 minutes, permeabilized with 0.2% Triton X-100 in PBS for 10 minutes at 4°C and blocked with PBS + 1% BSA for at least 1 hour. Primary antibodies used were: VRK1 (1:500, Sigma-Aldrich), Nestin rabbit (1:1000 Millipore), Nestin mouse (1:1000 R&D Systems), NCAM1 (1:300, Millipore), αSMA (1:400, Sigma-Aldrich), DDC (1:200, Abcam), Ki67 rabbit (1:200 Thermo Scientific) and SOX2 (1:500, Santa Cruz Biotech). Appropriate Alexa Fluor-conjugated secondary antibodies (1:1000, Invitrogen) were used, and nuclei were counterstained with DAPI (1 μg/ml). Actin was stained with Alexa Fluor 488 or 568 phalloidin (1:500) (Life Technologies). Images were acquired using a fluorescence microscope (Olympus BX61) and quantified using ImageJ software (NIH).

### Quantitative RT-PCR

RNA was extracted using the QIAamp RNeasy Mini Kit (Qiagen). For reverse transcription, 1 μg of RNA was converted to cDNA using qScript (QuantaBio). For RT-qPCR, Fast Sybr Green polymerase (Applied Biosystems) was used with glyceraldehyde-3-phosphate dehydrogenase (GAPDH) used as the control. Primers (0.2 μM, see Supplementary Table S1) and 50-300 μg of genetic material were used in a 7500 Fast Real Time PCR System (Applied Biosystems) in 96-well plates. Data analysis was performed using the ΔCt (cycle threshold) comparative method.

### Tumor xenografts

6–8 weeks-old CB-17 SCID mice were acquired from Harlan Laboratories. Each mouse was injected in the right flank with 4×10^5^ SK-N-SH cells, 24 h after transfection with the corresponding siRNAs. After 8 weeks, all mice were sacrificed humanely and tumors excised and measured. Tissue samples were included in paraffin for immunohistochemical analysis.

The generation and maintenance of PDXs was described previously (25). PDX tumor samples were obtained and treated similarly to xenograft.

### Statistical analysis

Data are presented as mean ± standard error of the mean (SEM) from at least three independent experiments. Statistical comparisons were performed using unpaired Student’s t-test for two groups or one-way ANOVA for multiple groups using GraphPad Prism 8.0. Fisher’s exact test was used for the analysis of positive and negative tumor samples. Correlation analyses used Pearson correlation coefficient. P-values<0.05 were considered statistically significant (*p<0.05, **p<0.01, ***p<0.001). In gene expression analysis plots, box and whiskers graphs show median, 10–90 percentiles and outliers as dots. The number of measurements or samples (n) is indicated in each case.

## Results

### VRK1 expression associates with an undifferentiated phenotype in neuroblastoma tumors

Even though VRK1 has been associated with the control of proliferation in various cancers (11), neuroblastoma cells show a widespread selective dependency to VRK1 expression compared with other solid tumor cells (mean NB selectivity: −0.847; mean other solid tumors selectivity: −0.393; NB cell line penetrance: 82%), demonstrating the importance of VRK1 function for NB progression (Supplementary Fig. S1 and Supplementary Table S2). Given the association of VRK1 with differentiation, development and stem cells in other tissues, we decided to analyze the expression of VRK1 in neuroblastoma tumors according to their differentiation status. VRK1 protein and mRNA are highly expressed on tumors with undifferentiated histology, compared to tumors with differentiated histology or rich in schwannian stroma (Fig. 1a-c). This association is maintained when restricted to MYCN non-amplified cases, indicating that the VRK1-differentiation relationship is independent of MYCN status (Fig. 1d). *VRK1* expression is increased in stage 4 tumors and inversely correlates with the expression of stablished differentiation markers like *DDC*, *NCAM1*, o *S100β* (Fig. 1e). Exploration of published single cell transcriptomic data (27,28) confirms the increased expression of *VRK1* in undifferentiated cell developmental stages of neuroblastoma, like the Schwan cell precursor cell (SPCs) population or the neural crest progenitor population (Supplementary Figure S2a-c). In summary, VRK1 expression is associated with an undifferentiated malignant phenotype in neuroblastoma tumors.

**Figure 1.**
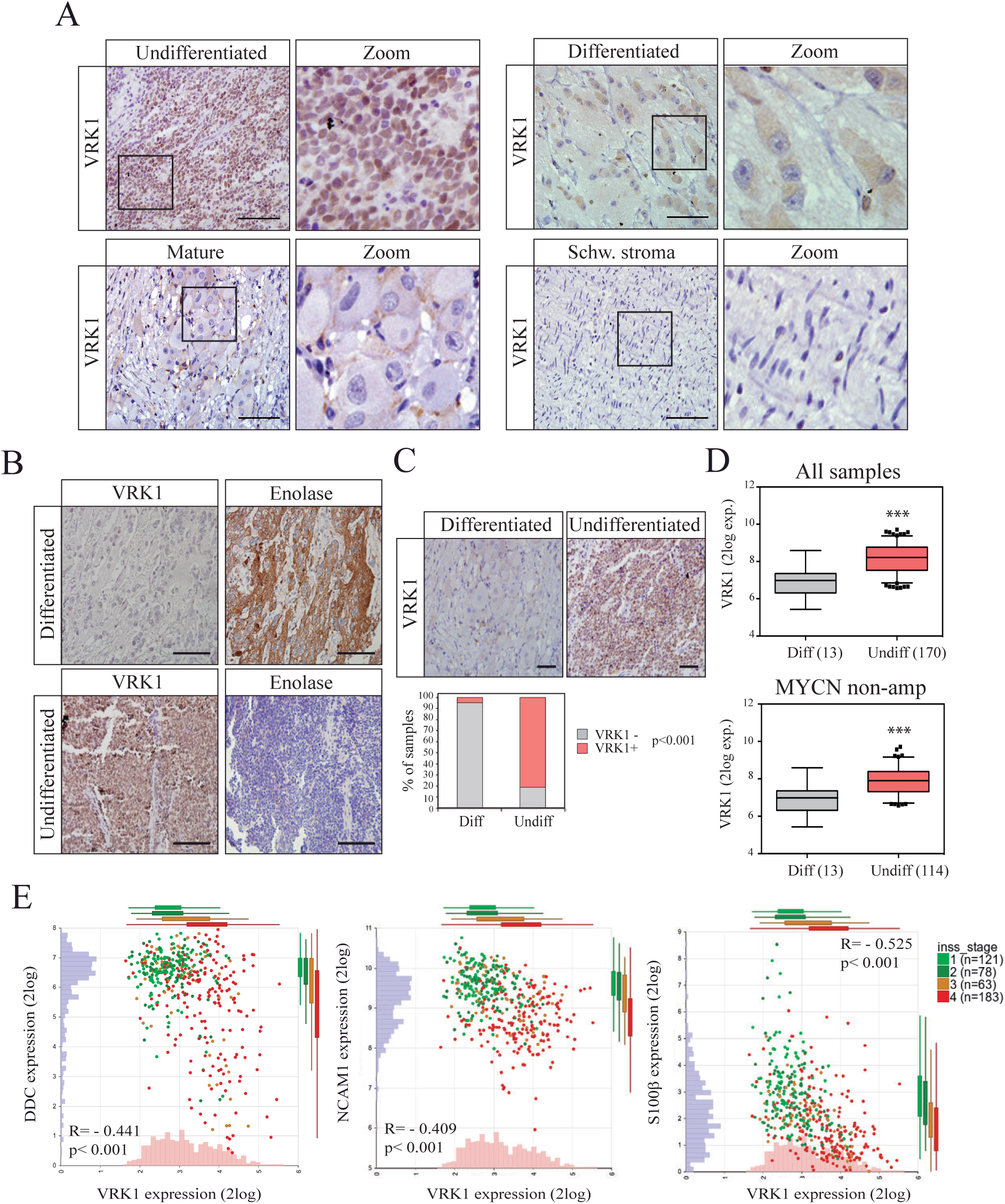
VRK1 expression inversely correlates with differentiation status. **A** Expression of VRK1 protein by immunohistochemistry in human neuroblastoma tumors with different histology. Zoom shows magnification from the indicated area. Bar: 100μm. **B** Representative immunohistochemical comparison between differentiated and undifferentiated tumor samples showing VRK1 protein and the differentiation marker enolase. Bar: 220μm. **C** VRK1 protein expression across human tumors in our TMA collection according to tumor histology. Quantification is shown. p<0.001, Fisher’s exact test. Bar: 100μm. **D** VRK1 gene expression according to tumor histology in NB patient tumors from the cohort GSE49710, or a MYCN non amplified subset. Number of tumors shown in brackets. ***: p<0.001. **E** Scatterplots showing tumor samples arranged according to VRK1 expression and the expression of the differentiation markers indicated in each graph. Individual tumor samples are colored according to INSS stage. Gene expression histograms and boxplots showing expression of each gene by stage are also shown. Number of tumors in each stage in brackets. Tumor cohort GSE45547.

### VRK1 inhibits differentiation and maintains an undifferentiated proliferative state in NB tumor cells

To further explore the association of VRK1 with differentiation we next induced the differentiation in culture of neuroblastoma cells. All-trans-retinoic acid treatment was used to force the neuronal differentiation in SK-N-SH and IMR-32 neuroblastoma cells, and TGFβ-1, to induce mesenchymal differentiation in the NB cell line SK-N-SH, not fully committed to neuronal lineage (Fig. 2a, b). We demonstrate effective differentiation by an increased expression of neuronal or mesenchymal markers and/or extensive neurite outgrowth. A marked reduction in VRK1 expression after differentiation was observed in all cases, indicating that VRK1 downregulation accompanies differentiation toward either lineage, a feature also observed in in vivo xenografts treated with retinoic acid (Supplementary Figure S2d).

**Figure 2.**
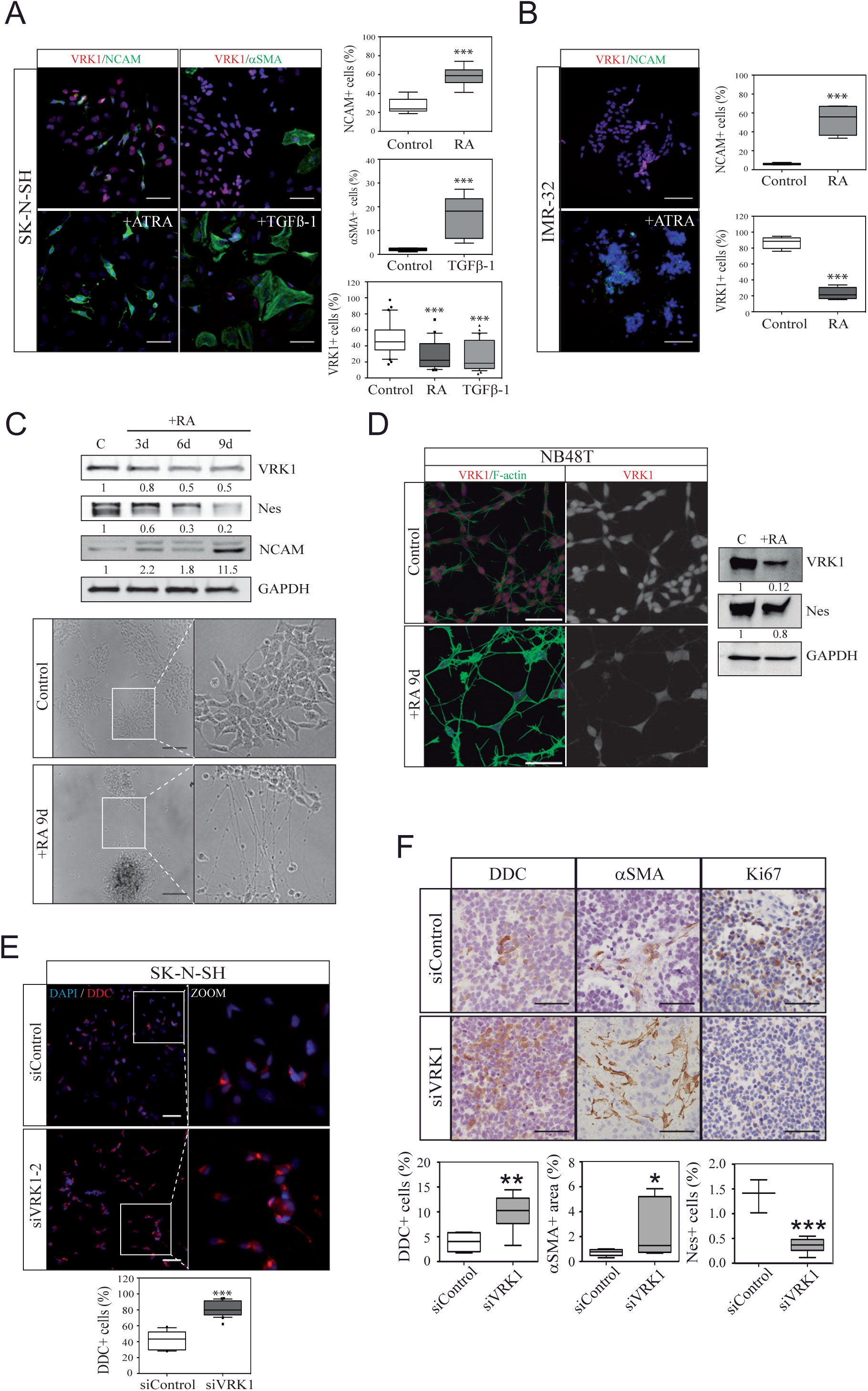
VRK1 inhibits differentiation in neuroblastoma cells. **A** Images showing labelling with VRK1, and the differentiation markers NCAM or αSMA in SK-N-SH NB cells after differentiation treatment with either all-trans retinoic acid (RA) or TFGβ1 (transforming growth factor beta-1). Quantifications are shown. Scale bars: 50 μm. **B** Images showing labelling with VRK1, and the differentiation marker NCAM in IMR-32 NB cells after differentiation with all-trans retinoic acid. Quantifications are shown. Scale bars: 50 μm. **C** Western blot showing the expression of the indicated markers at different time points after incubation with all-trans retinoic acid. Brightfield images show representative images of SK-N-SH cells in control condition or after 9 days incubation with retinoic acid. Note the presence of neurites in the differentiated condition. Scale bar:150 μm. **D** Images showing labelling with VRK1 and F-actin in the primary PDX-derived cell line NB48T after differentiation with all-trans retinoic acid. Scale bars: 50 μm. The western blot shows the expression of VRK1 and the undifferentiation marker Nestin before and after treatment. **E** Representative images and quantification showing increased DDC expression (red) in SK-N-SH cells transfected with siRNA against VRK1 compared to control. Scale bars: 100 μm. **F** Immunohistochemical detection of the differentiation markers DDC, αSMA or the proliferation marker Ki67 in tissue sections from xenografts obtained after injection of SK-N-SH cells or SK-N-SH cells treated with siRNA against VRK1. Scale bar: 60μm. Quantification of positive cells for the indicated differentiation markers in the tissue images are shown below. *: p<0.05; **: p<0.01; ***: p<0.001.

The downregulation of VRK1 expression was accompanied by a drop in the expression of the undifferentiation neural progenitor marker nestin (Fig. 2c). Similar changes were observed in the PDX-derived primary cell line NB48T (Fig. 2d).

To determine whether VRK1 actively maintains the undifferentiated state, we performed loss-of-function experiments using siRNA-mediated knockdown. VRK1 downregulation alone triggered neuronal differentiation in SK-N-SH cells (Fig. 2e). In tumor xenografts formed by cells treated with siRNA against VRK1, an increase in the expression of neuronal and mesenchymal differentiation markers, and a decrease of nestin were observed, indicating a long-lasting effect in vivo after the manipulation of VRK1 levels (Fig. 2f) (18). Ki67-positive proliferating cells were also decreased, demonstrating that VRK1 is required for maintaining high proliferative capacity. Taken together, these findings indicate that VRK1 functions as a master regulator that simultaneously maintains proliferative capacity and actively suppresses multiple differentiation pathways in neuroblastoma cells.

### VRK1 promotes NB proliferative progenitor cells maintenance

As VRK1 expression is generally associated with proliferation potential in neuroblastoma tumors, we set to discriminate VRK1 function in tumors with favorable (differentiated) histology versus unfavorable (undifferentiated) histology. To do that we identified genes whose expression correlate with a high expression of VRK1 in tumors with favorable or/and unfavorable histology. We next performed enrichment analysis identifying proliferation control and cell cycle signatures associated with VRK1 co-expression in all tumors, meanwhile specifics stemness/oncogenic signatures appeared mainly in tumors with unfavorable histology (Fig. 3a and Supplementary Table S3).

**Figure 3.**
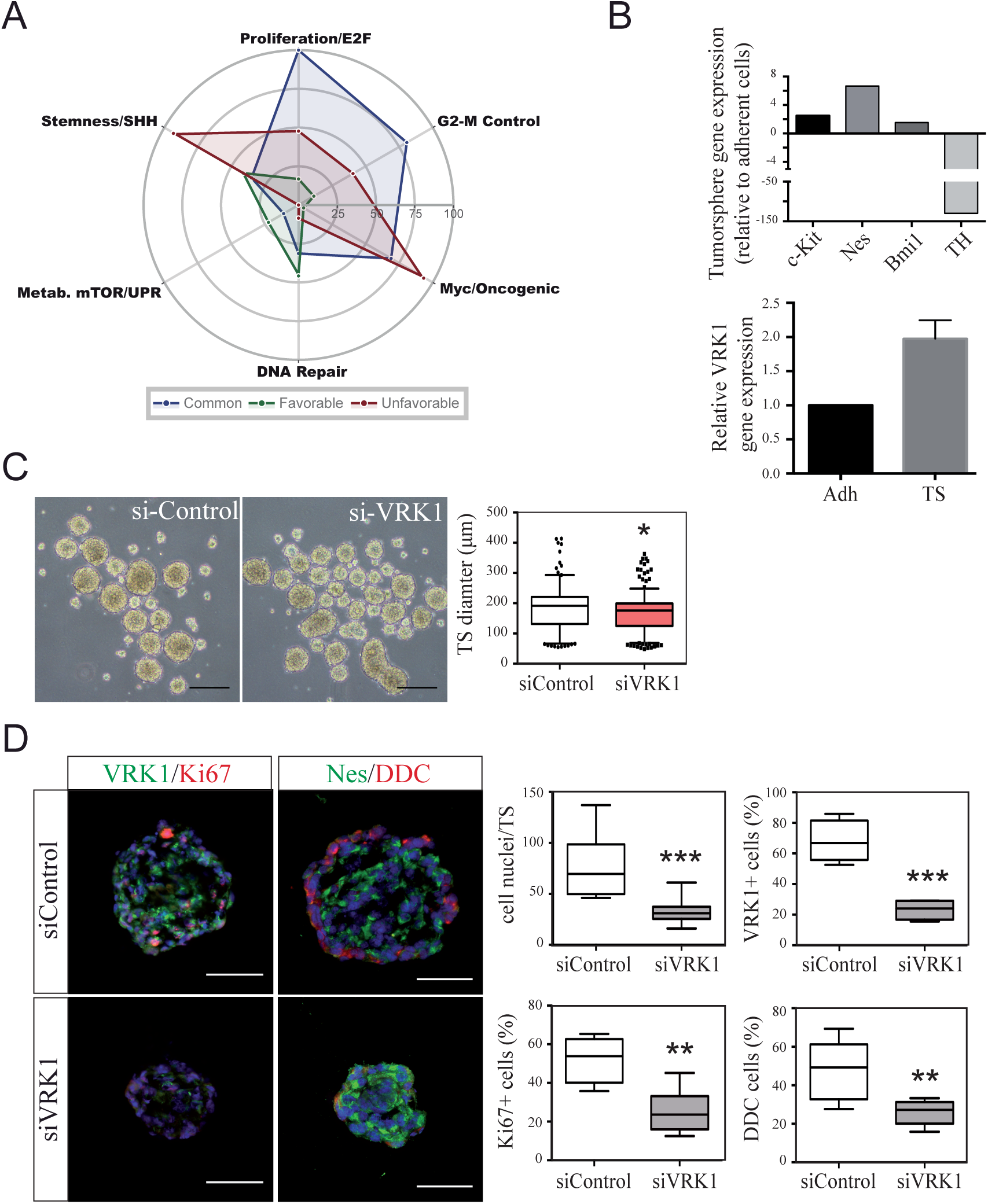
VRK1 is enriched in tumorspheres and required for stem cell-like properties. **A** Functional landscape of the VRK1 co-expression gene groups in neuroblastoma. Radar chart displaying normalized enrichment scores (0–100) across six functional axes for the Common, Favorable, and Unfavorable gene groups. Scores were calculated by normalizing the – log₁₀(adjusted p-value) of the most significant term per category against the global maximum observed (E2F Targets, Common group = 109.2). Databases used: MSigDB Hallmarks, MSigDB Oncogenic Signatures, GO Biological Process 2026, and CellMarker 2024. **B** qRT-PCR analysis comparing gene expression between adherent (Adh) and tumorsphere (TS) cultures from SK-N-SH cells showing enrichment of stem cell markers (*c-Kit*, *Nes*, *BMI1*, *VRK1*) and downregulation of differentiation marker (*TH*). **C** Representative images and quantification of tumorsphere formation with SK-N-SH cells after VRK1 knockdown or control. Scale bars: 200 μm. **D** Immunofluorescence of tumorspheres obtained after VRK1 knockdown or control showing the expression of differentiation markers. Scale bars: 100 μm. Data represent mean ± SEM from three independent experiments. *p<0.05, **p<0.01, ***p<0.001.

Undifferentiated neuroblastoma tumors are enriched in cells with neural-crest stem like features with malignant properties (6,25). To further investigate the role of VRK1 in NB differentiation, we explored the role of VRK1 in these cells using previously characterized tumorsphere cultures to select and expand stem-like cell populations (26). Gene expression analysis of undifferentiated tumorspheres compared to adherent cultures revealed upregulation of stem cell markers, downregulation of the neuronal differentiation marker TH, and notably, a significant increase in VRK1 expression (Fig. 3b), indicating that VRK1 is not merely ubiquitously expressed in proliferating cells but is preferentially enriched in stem-like, self-renewing subpopulations. Tumorspheres formed by VRK1 knocked-down NB cells show a significant decrease in diameter, related to their proliferative capacity (Fig. 3c). The obtained tumorspheres do not seem fully developed, showing a marked decrease in intra-sphere undifferentiated cell proliferation and loss of the characteristic peripheral DDC+, neuronal differentiated derivatives, indicating a role for VRK1 in stem cell self-renewal and function (Fig. 3d).

### VRK1 regulates the stemness transcription factor SOX2

To understand the mechanism by which VRK1 maintains stemness and prevents differentiation, we investigated its relationship with key transcriptional regulators of stem cell identity. Among them, *SOX2* is a core pluripotency transcription factor that maintains neural stem cells (29). *SOX2* is frequently expressed in neuroblastoma and has emerged as a potential downstream effector of VRK1 in glioblastoma stem cells and the skin stem cell compartment (19).

VRK1 is co-expressed with SOX2 in neuroblastoma tumors (Fig. 4a). VRK1 knock-down in undifferentiated tumorspheres-derived cells leads to a decrease of SOX2 protein expression in the nucleus (Fig. 4b-c). Single cell co-expression analysis shows a positive correlation between VRK1 and SOX2 protein expression levels, although some cells with low VRK1 expression can still maintain considerable expression of SOX2. Meanwhile, VRK1 overexpression leads to an increase in SOX2 levels (Fig. 4d).

**Figure 4.**
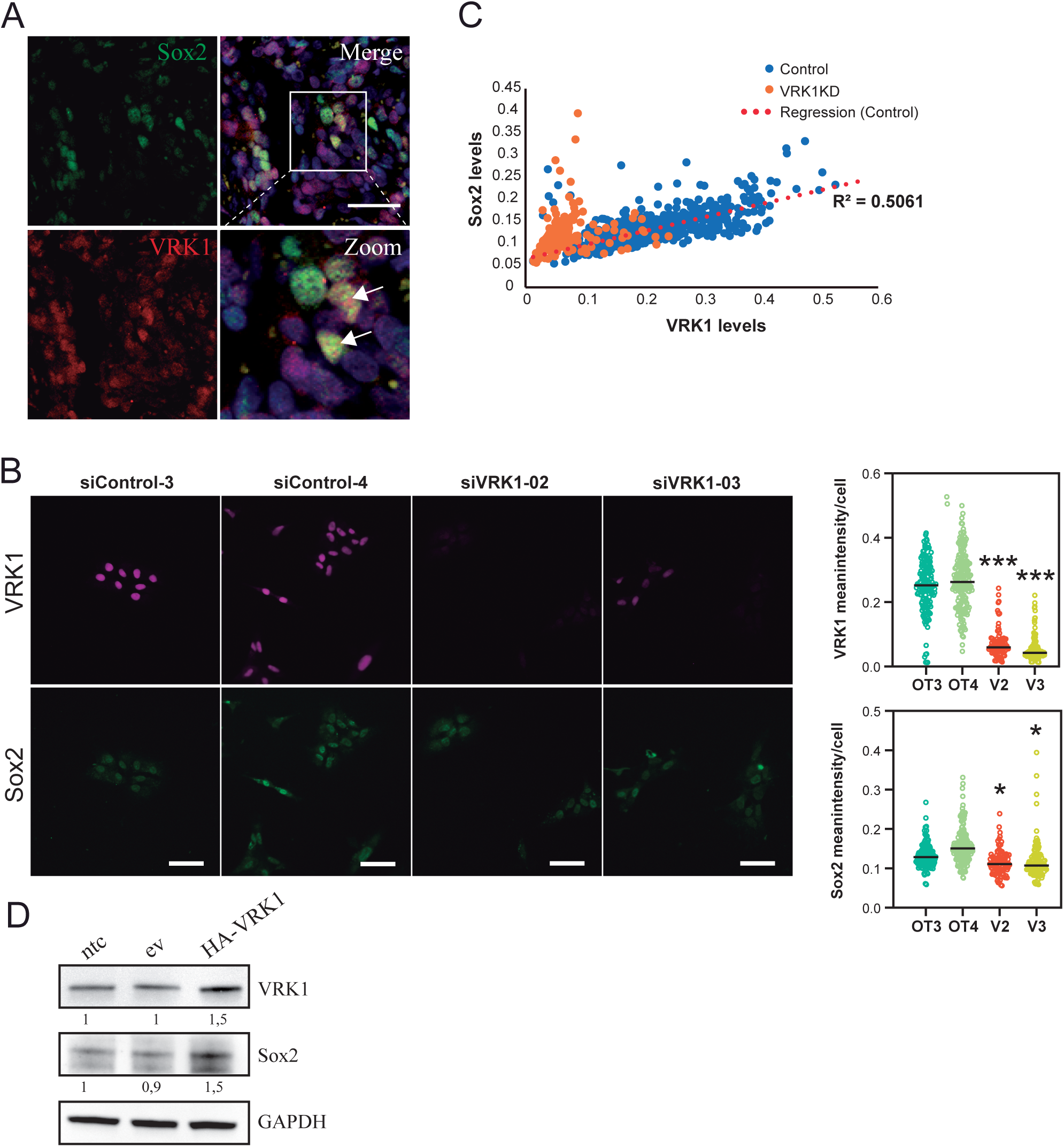
VRK1 regulates the stemness transcription factor SOX2. **A** Representative immunofluorescence images showing nuclear co-localization of VRK1 and SOX2 in a PDX tumor. Arrows indicate co-localizing nuclei. Scale bars: 50 μm. **B** Representative immunofluorescence images showing VRK1 and SOX2 staining in siControl or VRK1 knockdown treated tumorspheres-derived cells. Scale bars: 50 μm. Dot plots show distribution of VRK1 and SOX2 intensities across different conditions. **C** Single-cell quantitative analysis of fluorescence signal in tumorspheres-derived cells showing correlation between VRK1 and SOX2 intensities. **D** Western blot showing SOX2 levels after VRK1 overexpression (HA-VRK1) GAPDH serves as a loading control.

In agreement with published data in other cancer models, VRK1 directly or indirectly stabilizes the core stemness transcription factor *SOX2* in neuroblastoma cells. VRK1 has been shown to directly bind SOX2 and SOX2 to transcriptionally activate *VRK1* (19). Epigenetics analysis of the *VRK1* locus (chr14q23.2) through publicly available data in ENCODE revealed active chromatin marks (H3K4me3 and H3K27ac) consistently detected in SK-N-SH neuroblastoma cells, progenitor neural cells and neural crest cells (Supplementary Figure S3 and Supplementary Table S4), indicating an accessible chromatin state. JASPAR prediction identified multiple possible sites for SOX2 binding in the *VRK1* promoter region, 5 of them with a high confidence (relative score>=0.9). Moreover, and a search on the ReMap2022 DNA-binding sequencing repository identified 5 non-redundant peaks for SOX2 binding in the *VRK1* locus in several contexts, including embryonic stem cells and neural stem cells (Supplementary Figure S3), 13 of them coinciding with JASPAR predictive motives, although none in the promoter region. Interestingly, no picks for SOX9 or SOX10 binding were identified, revealing the possible specificity for SOX2 in regulating *VRK1* transcription.

In summary, VRK1 manipulation alters SOX2 levels, providing a mechanistic link between VRK1’s role in maintaining the undifferentiated, stem-like phenotype and the transcriptional control of stemness.

## Discussion

The data presented here converge on a model where VRK1 marks, and is functionally required to sustain the undifferentiated stem-like compartment in neuroblastoma. We previously described VRK1 general involvement in neuroblastoma tumor proliferation (18) and analysis of DepMap CRISPR dependency data demonstrates a NB selective requirement for VRK1. Our results show that VRK1 consistently emerges as a gatekeeper that suppresses differentiation and promotes stem-like traits and proliferation.

Gene correlation analysis with VRK1 in a NB tumor cohort stratified according to histology shows specific enrichment in signatures related to stemness programs in tumors with undifferentiated histology, meanwhile a general association of VRK1 with proliferation programs is detected in all tumors, independently of differentiation status. This points towards a possible role for VRK1 in differentiation control, besides the known functions in proliferation described previously for neuroblastoma and other tumors (11). VRK1 is expressed during human cortex development, and mutations in VRK1 are linked to severe neuronal phenotypes, including brain developmental defects and degeneration of spinal motor neurons, indicating its possible role in neurogenesis (30–32). The diverse nature of substrates described for VRK1 phosphorylation and VRK1 capacity for chromatin remodeling might indicate cell or context-specific functions controlling proliferation or differentiation.

VRK1 expression was significantly enriched in undifferentiated versus differentiated tumor samples, and VRK1 transcript levels were significantly higher in undifferentiated tumors both in the full cohort and after restricting the analysis to MYCN-non-amplified cases. The finding that VRK1 association with differentiation and aggressiveness persists even in MYCN non-amplified undifferentiated cases suggests that VRK1 operates through a pathway independent of or parallel to MYCN, potentially representing a broader mechanism of differentiation control, and is consistent with our earlier report that VRK1 predicts malignancy independently of MYCN (18). Our data suggests VRK1 could serve as a robust biomarker for neuroblastoma differentiation, indicating a diagnostic utility.

Our results with pro-differentiation agents, show that VRK1 loss follows pharmacological differentiation in NB cells, independently of the lineage. Genetically silencing VRK1 expression reduces proliferation/stemness marker expression, coupled to a reciprocal rise in differentiation markers, indicating that VRK1 is not merely a passive marker of the undifferentiated state but is functionally required to restrain differentiation. VRK1 loss is thus shown as a potent inducer of differentiation programs, suggesting it sits at a central node repressing diverse differentiation pathways.

Concordantly with previously described associations in other contexts (19,21,22) and the gene enrichment data in neuroblastoma tumors, VRK1 was upregulated in stem-like tumorspheres cells alongside known stem/progenitor markers. VRK1 also tracks proliferative, lineage-uncommitted cell states in the sympathoadrenal lineage from which neuroblastoma arises. This demonstrates that VRK1 is not merely ubiquitously expressed in proliferating cells but is preferentially enriched in stem-like, progenitor subpopulations. Our results show that VRK1 is critical for neuroblastoma cancer stem cell–like properties, supporting undifferentiated tumorsphere formation, proliferation, and expression of key stemness markers. Its depletion undermines self-renewal and reduces sphere growth. Conversely to what we have described in the bulk cell culture, tumorspheres show a reduction of DDC derivatives after VRK1 knock-down, probably the results of severely compromising stem cells viability and function.

Among the core pluripotency regulators with a role in neural crest stem cells specification, SOX (SRY-box) transcription factors, mainly SOX2, SOX9 and SOX10, emerge as main players with potential functions during neuroblastoma biology (29,33,34). SOX2 has been previously associated with VRK1 in other contexts, establishing reciprocal regulation that sustain the proliferative undifferentiated state during tissue regeneration and in tumors (19). In the context of neuroblastoma tumors and cell lines, we have established a positive correlation between VRK1 and SOX2 expressions. Gain-of-function experiments suggest a VRK1-dependent regulation of SOX2, although a reciprocal SOX2-dependent regulation of VRK1 or a shared upstream driver cannot be ruled out. Previous reports demonstrated that SOX2 transcriptionally upregulates VRK1 via its proximal promoter, and we have shown active chromatin state and potential SOX2 binding in neuroblastoma related contexts. All together, these results raise the possibility of a mutually reinforcing VRK1-SOX2 relationship that helps lock neuroblastoma cells into an undifferentiated state, stabilizing a core transcriptional program that underlies neuroblastoma stemness. Although a direct binding between VRK1 and SOX2 has been described in breast carcinoma and teratoma cells, and phosphorylation suggested in in-vitro assays, the mechanism by which VRK1 and SOX2 interact in neuroblastoma cells remains to be identified and is a limitation of the present study. Candidate indirect mediators could include shared upstream regulators within the neural crest/stemness identity module, or an intermediate signaling step downstream of VRK1 kinase activity. For example, the AP-1 transcription factors c-JUN and FOS are known to be phosphorylated by VRK1 (35) and are part of SOX2 signaling in neural stem cells (36).

We propose a model in which, SOX2-high neuroblastoma cells induce VRK1 as part of a stem-like proliferative program, VRK1 supports expansion of that state and actively blocks differentiation. Both markers decrease as cells move toward neural differentiation. Targeting VRK1 could enforce differentiation across NB subtypes and simultaneously curb proliferation, offering a dual mechanism to reduce tumor aggressiveness.

## Acknowledgements

We thank Dr Pedro A. Lazo, from Centro de Investigación del Cáncer in Salamanca, for kindly providing reagents and helpful discussion. We are grateful to the Andalusian tissue Biobank at Virgen del Rocio University Hospital for their help with the human tumor samples and tissue histology. We thank Dr. Catalina Marquez, Dr. Gema Ramirez and Dr. Rosa Cabello from the Virgen del Rocio University Hospital for their insight and help obtaining NB samples.

## Authors’ contributions

FMV and RP designed and supervised the study. FMV wrote the manuscript. FMV, MO, AC and MAG developed methodology. FMV, MO, MAG, AC and AA acquired data. IR provided technical support and FMV, MO, AC, MAG and RP analyzed and interpreted the data. All authors approved the final manuscript.

## Availability of data and materials

The datasets supporting the conclusions of this article are included within the article and its supplementary files.

## Supplementary Figure legends

**Figure S1**. VRK1 dependency in NB cells. **A** Distribution of NB VRK1 CRISPR KO gene effect in different NB cell lines or all other solid tumor cell lines. **B** Plot showing the mean dependency of different genes in NB cell lines versus all other solid tumor cell lines. Key neuroblastoma driver genes are highlighted. **C** Plot showing the effect size (dependency in NB cell lines-dependency in all other cancers) for different genes versus the fraction of dependent NB cell lines for those genes. Key neuroblastoma driver genes are highlighted. Data from DepMap Public 25Q3+Score. Minimum effect size = 0.25. Minimum fraction= 0.2. p value<0.05. Full data for top dependency genes can be seen on Supplementary Table S2.

**Figure S2. VRK1 expression in NB developmental stages. A** Heatmap showing the expression in different developmental neuroblastoma cell state of the listed genes according to (27). Neural crest progenitor (NC_Prog), Bridge-Schwann cell precursor cells (Bridge-SCP), mesenchymal differentiation (Diff_MES), neuroblastic adrenergic differentiation (Diff_Neuroblast), and chromaffin cell states are shown. **B** UMAP showing annotation of the different cell types in human adrenal medulla according to (28). **C** Expression of *VRK1* in the annotated clusters in B. UMAP and violin plots are shown. **D** Expression of *VRK1* in PDX-derived cells treated with retinoic acid or the corresponding controls. Dots represent single data points and are colored according to CD44 expression, a marker of undifferentiated NB cells. Data from GSE120920.

**Figure S3. Epigenetic regulation of the VRK1 gene.** The diagram shows a schematic representation of the human VRK1 gene (chr14:96,797,382–96,881,609; GRCh38/hg38), confirmed on the plus (+) strand (Ensembl ENSG00000100749). The 5kb proximal promoter, 5’ and 3’ UTRs and VRK1 CDS are indicated. Published SOX2 binding ChIP-seq peaks from ReMap2022 by cell-type biotype (hiPSC, Tyroid tumors, RENVM, HNSC and HCC95) spanning the VRK1 locus are shown side by side, drawn to scale as a positional reference. They are shown aligned with JASPAR-predicted SOX2 binding motifs (matrix MA0143.4). Jaspara sites overlapping ReMap peaks and high-confidence JASPAR motif (relative score ≥0.90) are highlighted. Below, the chromatin landscape of the VRK1 locus is shown on relevant Chip-seq datasets revealing two active-chromatin histone marks, H3K4me3 (canonical promoter mark) and H3K27ac (active promoter/enhancer mark) and including sources relevant to neurodevelopmental and neuroblastoma biology (human neural stem cells, human neural crest and the SK-N-SH neuroblastoma cell line). ENCODE experiment codes shown on the right. Peaks with signalValue ≥20 are marked.

**Supplementary Table S1:** Oligonucleotide primers sequence used in the study.

**Supplementary Table S2:** Context dependency DepMap results for neuroblastoma cell lines.

**Supplementary Table S3:** VRK1 correlated genes in tumors according to histology.

**Supplementary Table S4:** Data epigenetics regulation VRK1 gene by Sox2.

## References

1. Sainero-Alcolado L, Sjöberg Bexelius T, Santopolo G, Yuan Y, Liaño-Pons J, Arsenian-Henriksson M. Defining neuroblastoma: From origin to precision medicine. Neuro Oncol.; 2024;26:2174–92.

2. Johnsen JI, Dyberg C, Wickström M. Neuroblastoma-A Neural Crest Derived Embryonal Malignancy. Front Mol Neurosci. 2019;12:9.

3. Gomez RL, Ibragimova S, Ramachandran R, Philpott A, Ali FR. Tumoral heterogeneity in neuroblastoma. Biochimica et Biophysica Acta (BBA) - Reviews on Cancer. 2022;1877:188805.

4. Zeineldin M, Patel AG, Dyer MA. Neuroblastoma: When differentiation goes awry. Neuron. Neuron; 2022;110:2916–28.

5. He G, He S, Jing X, Dai Y, Guo X, Gao J, et al. Dissecting neuroblastoma heterogeneity through single-cell multi-omics: insights into development, immunity, and therapeutic resistance. Oncogene. 2025;1–17.

6. Bedoya-Reina OC, Li W, Arceo M, Plescher M, Bullova P, Pui H, et al. Single-nuclei transcriptomes from human adrenal gland reveal distinct cellular identities of low and high-risk neuroblastoma tumors. Nat Commun. 2021;12:5309.

7. Grossmann LD, Chen C-H, Uzun Y, Thadi A, Wolpaw AJ, Louault K, et al. Identification and Characterization of Chemotherapy-Resistant High-Risk Neuroblastoma Persister Cells. Cancer Discov. 2024;14:2387–406.

8. Valbuena A, Sanz-García M, López-Sánchez I, Vega FM, Lazo PA. Roles of VRK1 as a new player in the control of biological processes required for cell division. Cell Signal. 2011;23:1267–72.

9. Valbuena A, López-Sánchez I, Lazo PA. Human VRK1 Is an Early Response Gene and Its Loss Causes a Block in Cell Cycle Progression. Williams S, editor. PLoS One. 2008;3:e1642.

10. So J, Mabe NW, Englinger B, Chow K-H, Moyer SM, Yerrum S, et al. VRK1 as a synthetic lethal target in VRK2-methylated cancers of the nervous system. JCI Insight. American Society for Clinical Investigation; 2022; Oct 10;7(19):e158755

11. Campillo-Marcos I, García-González R, Navarro-Carrasco E, Lazo PA. The human VRK1 chromatin kinase in cancer biology. Cancer Lett. Cancer Lett; 2021;503:117–28.

12. Liu Z-C, Cao K, Xiao Z-H, Qiao L, Wang X-Q, Shang B, et al. VRK1 promotes cisplatin resistance by up-regulating c-MYC via c-Jun activation and serves as a therapeutic target in esophageal squamous cell carcinoma. Oncotarget. 2017;8:65642–58.

13. Mon AM, MacKinnon AC, Traktman P. Overexpression of the VRK1 kinase, which is associated with breast cancer, induces a mesenchymal to epithelial transition in mammary epithelial cells. Ahmad A, editor. PLoS One. 2018;13:e0203397.

14. Santos CR. VRK1 Signaling Pathway in the Context of the Proliferation Phenotype in Head and Neck Squamous Cell Carcinoma. Molecular Cancer Research. 2006;4:177–85.

15. Campillo-Marcos I, Lazo PA. Implication of the VRK1 chromatin kinase in the signaling responses to DNA damage: a therapeutic target? Cell Mol Life Sci.; 2018;75:2375–88.

16. Huang W, Cui X, Chen Y, Shao M, Shao X, Shen Y, et al. High VRK1 expression contributes to cell proliferation and survival in hepatocellular carcinoma. Pathol Res Pract. 2016;212:171–8.

17. Shields JA, Meier SR, Bandi M, Mulkearns-Hubert EE, Hajdari N, Ferdinez MD, et al. VRK1 Is a Synthetic-Lethal Target in VRK2-Deficient Glioblastoma. Cancer Res.; 2022;82:4044–57.

18. Colmenero-Repiso A, Gómez-Muñoz MA, Rodríguez-Prieto I, Amador-Álvarez A, Henrich K-O, Pascual-Vaca D, et al. Identification of VRK1 as a New Neuroblastoma Tumor Progression Marker Regulating Cell Proliferation. Cancers. 2020;12:3465.

19. Moura DS, Fernández IF, Marín-Royo G, López-Sánchez I, Martín-Doncel E, Vega FM, et al. Oncogenic Sox2 regulates and cooperates with VRK1 in cell cycle progression and differentiation. Sci Rep. 2016;6:28532.

20. Vega FM, Gonzalo P, Gaspar ML, Lazo PA. Expression of the VRK (vaccinia-related kinase) gene family of p53 regulators in murine hematopoietic development. FEBS Lett. 2003/06/05. 2003;544:176–80.

21. Choi YH, Park C-H, Kim W, Ling H, Kang A, Chang MW, et al. Vaccinia-Related Kinase 1 Is Required for the Maintenance of Undifferentiated Spermatogonia in Mouse Male Germ Cells. Milstone DS, editor. PLoS One; 2010;5:e15254.

22. Wen Y, Zhou S, Gui Y, Li Z, Yin L, Xu W, et al. hnRNPU is required for spermatogonial stem cell pool establishment in mice. Cell Rep. Cell Rep; 2024;43.

23. Bankhead P, Loughrey MB, Fernández JA, Dombrowski Y, McArt DG, Dunne PD, et al. QuPath: Open source software for digital pathology image analysis. Sci Rep. 2017;7:16878.

24. Allred DC, Harvey JM, Berardo M, Clark GM. Prognostic and predictive factors in breast cancer by immunohistochemical analysis. Modern Pathology. 1998. page 155–68.

25. Vega FM, Colmenero-Repiso A, Gómez-Muñoz MA, Rodríguez-Prieto I, Aguilar-Morante D, Ramírez G, et al. CD44-high neural crest stem-like cells are associated with tumour aggressiveness and poor survival in neuroblastoma tumours. EBioMedicine. 2019;49:82–95.

26. Amador-Álvarez A, Gómez-Muñoz MA, Rodríguez-Prieto I, Pardal R, Vega FM. A protocol to enrich in undifferentiated cells from neuroblastoma tumor tissue samples and cell lines. STAR Protoc. Cell Press; 2022;3:101260.

27. Xu Y, Lou D, Chen P, Li G, Usoskin D, Pan J, et al. Single-cell MultiOmics and spatial transcriptomics demonstrate neuroblastoma developmental plasticity. Dev Cell.; 2025;60:2248–2263.e11.

28. Jansky S, Sharma AK, Körber V, Quintero A, Toprak UH, Wecht EM, et al. Single-cell transcriptomic analyses provide insights into the developmental origins of neuroblastoma. Nat Genet. Nature Publishing Group; 2021;53:683–93.

29. Tsaytler P, Blaess G, Scholze-Wittler M, Koch F, Herrmann BG. Early neural specification of stem cells is mediated by a set of SOX2-dependent neural-associated enhancers. Stem Cell Reports. Cell Press; 2024;19:618–28.

30. Apridita Sebastian W, Shiraishi H, Shimizu N, Umeda R, Lai S, Ikeuchi M, et al. Ankle2 deficiency-associated microcephaly and spermatogenesis defects in zebrafish are alleviated by heterozygous deletion of vrk1. Biochem Biophys Res Commun.; 2022;624:95–101.

31. Vinograd-Byk H, Renbaum P, Levy-Lahad E. Vrk1 partial Knockdown in Mice Results in Reduced Brain Weight and Mild Motor Dysfunction, and Indicates Neuronal VRK1 Target Pathways. Sci Rep. 2018;8:11265.

32. de Albuquerque Bueno MG, Dos Santos DF, Rossor AM, Laura M, Horga A, Frezatti RSS, et al. VRK1-Related Motor Neuropathy With Upper Motor Neuron Signs and Selective Muscle Involvement. J Peripher Nerv Syst.; 2026;31:e70145.

33. Nagashimada M, Ohta H, Li C, Nakao K, Uesaka T, Brunet J-F, et al. Autonomic neurocristopathy-associated mutations in PHOX2B dysregulate Sox10 expression. Journal of Clinical Investigation. 2012;122:3145–58.

34. Sarkar A, Hochedlinger K. The sox family of transcription factors: versatile regulators of stem and progenitor cell fate. Cell Stem Cell. 2013/01/08. 2013;12:15–30.

35. Sevilla A, Santos CR, Barcia R, Vega FM, Lazo PA. c-Jun phosphorylation by the human vaccinia-related kinase 1 (VRK1) and its cooperation with the N-terminal kinase of c-Jun (JNK). Oncogene. 2004;23:8950–8.

36. Pagin M, Pernebrink M, Giubbolini S, Barone C, Sambruni G, Zhu Y, et al. Sox2 controls neural stem cell self-renewal through a Fos-centered gene regulatory network. Stem Cells; 2021;39:1107–19.

