## Supplementary figures and images for "VRK1 kinase maintains an undifferentiated proliferative state in neuroblastoma tumor cells"

### Supplementary Figure S1

#### Figure S1

A

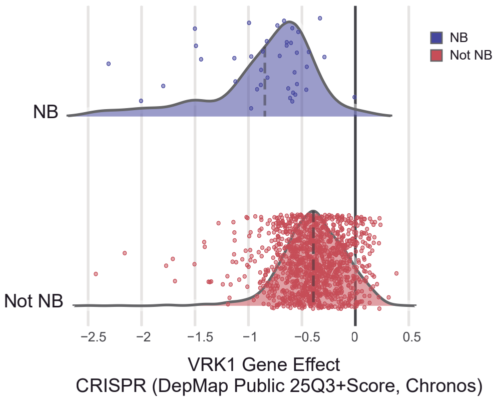


B

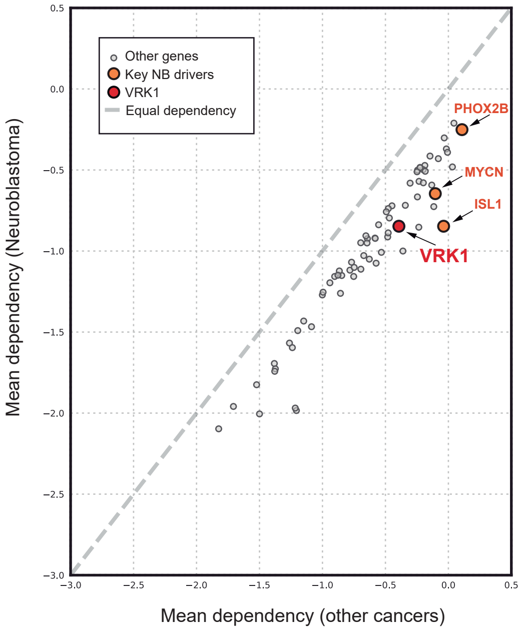


C

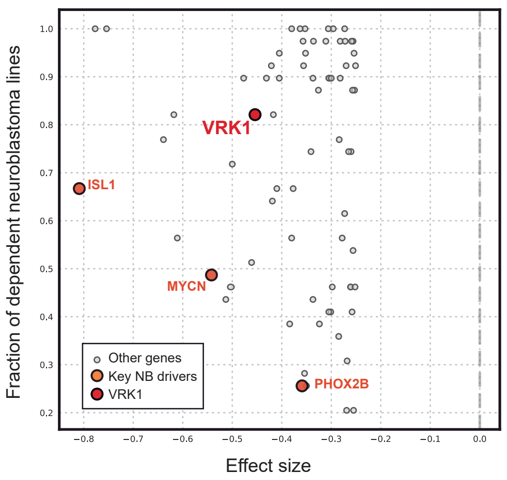

### Supplementary Figure S2

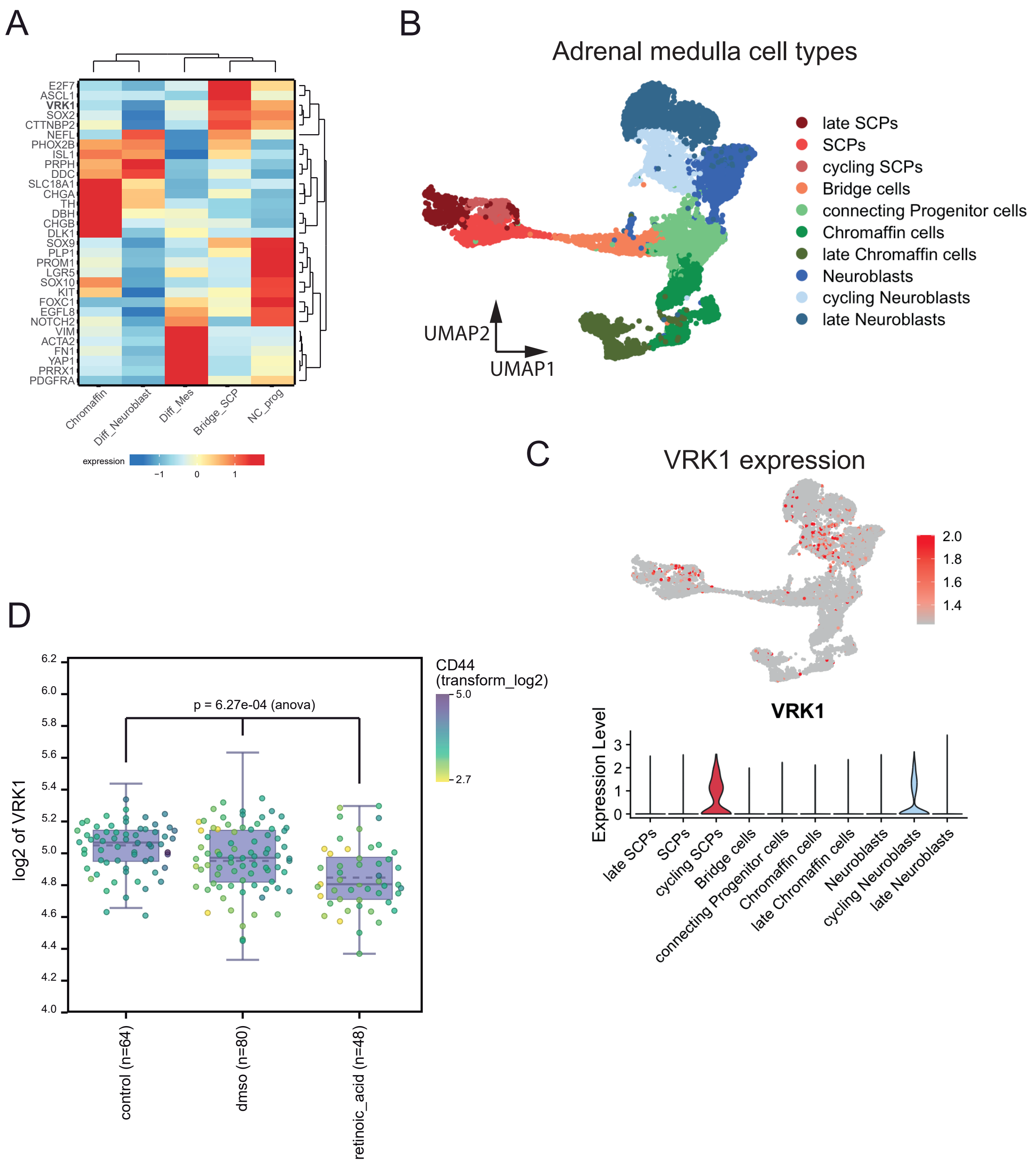

### Supplementary Figure S3

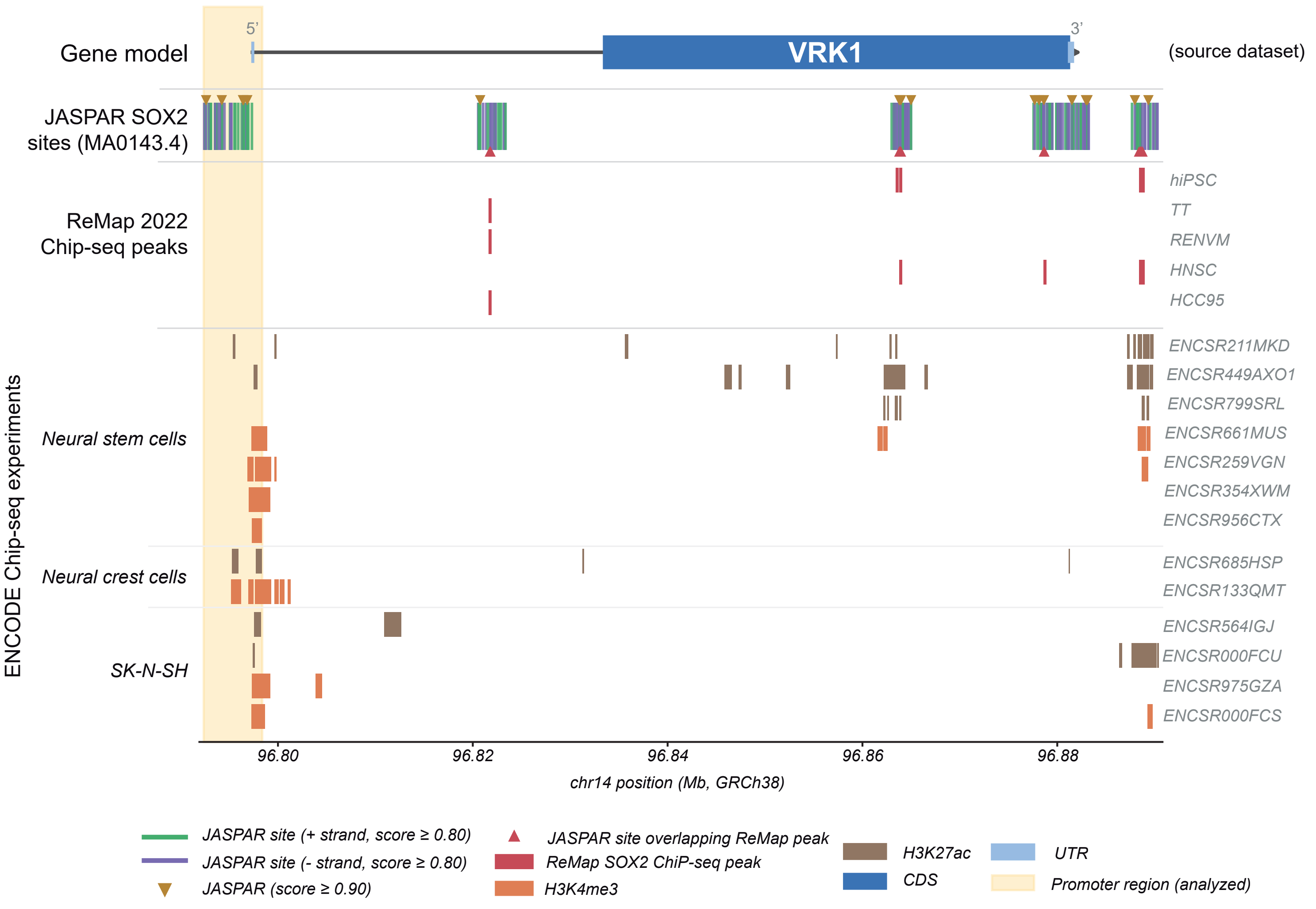
